# Proteins required for vacuolar function are targets of lysine polyphosphorylation in yeast

**DOI:** 10.1101/697847

**Authors:** Liam McCarthy, Amanda Bentley-DeSousa, Alix Denoncourt, Yi-Chieh Tseng, Matthew Gabriel, Michael Downey

**Affiliations:** Department of Cellular and Molecular Medicine, University of Ottawa, Ottawa, Ontario, Canada, K1H 8M5; Ottawa Institute of Systems Biology, University of Ottawa, Ottawa, Ontario, Canada, K1H 8M5

**Author notes:** Contributed equally to this work (Liam McCarthy, Amanda Bentley-DeSousa).

## Abstract

Polyphosphates (polyP) are long chains of inorganic phosphates that can be attached to lysine residues of target proteins as a non-enzymatic post-translational modification. This modification, termed polyphosphorylation, may be particularly prevalent in bacterial and fungal species that synthesize and store large quantities of polyP. In this study, we applied a proven screening strategy to evaluate the polyphosphorylation status of over 200 candidate targets in the budding yeast *S. cerevisiae.* We report 8 new polyphosphorylated proteins that interact genetically and physically with a previously identified network of targets implicated in ribosome biogenesis. The expanded target network includes vacuolar proteins Prb1 and Apl5, whose modification with polyP suggests a model for feedback regulation of polyP synthesis, while raising additional questions regarding the location of polyphosphorylation *in vivo.*

## INTRODUCTION

Polyphosphate (polyP) is a polymer of inorganic phosphate moieties joined together in linear chains ranging from 3-1000s of residues in length. Evidence suggests that these chains exist in all cells, although their length and concentration vary widely. PolyP is particularly abundant in the budding yeast *S. cerevisiae*, comprising upwards of 10 % of the dry weight of the cell and reaching internal concentrations of over 200 mM in terms of individual phosphate units (1,2). Most of the polyP in yeast is stored in the vacuole following synthesis by the vacuolar membrane-bound VTC complex (3). Overall chain length is controlled by exo- and endophosphatases Ddp1, Ppn1, Ppn2, and Ppx1 (4). In addition to its role in ion homeostasis and phosphate metabolism (5,6), Azevedo *et al.* showed that polyP chains can be non-covalently added to lysine residues as a post-translational modification (PTM) (7). They characterized this modification on proteins Nsr1 and Top1, providing evidence that polyP chains can regulate protein interactions and topoisomerase activity of purified Top1 *in vitro*, as well as the localization of Nrs1 and Top1 *in vivo*. Polyphosphorylation is non-enzymatic and occurs in poly-acidic serine and lysine rich (PASK) motifs (7,8). While the sequence requirements for polyP addition are unclear, lysine residues appear to be the modified site (7,9,10). Consistent with this, mutation of lysine residues within predicted PASK motifs to arginine prevents polyphosphorylation of Nsr1, Top1, and Rpa34 (7,9). Polyphosphorylation causes dramatic electrophoretic shifts when protein extracts are run on denaturing Bis-Tris NuPAGE gels, but this same shift does not occur with traditional SDS-PAGE gels (7,9). The underlying reasons for this difference are not completely clear, although it could be related to the use of TEMED in the polymerization of SDS-PAGE gels (9).

Recently, we reported the identification of an additional 15 polyphosphorylation targets in yeast, including a conserved network implicated in ribosome biogenesis (9). Many of the PASK motifs in these proteins are contained within intrinsically disordered regions that appear to be under evolutionary selection (11). Based on enrichment in ribosome-related functions, we uncovered a novel function for *VTC4*, encoding the catalytic subunit of the VTC complex (12), in polysome assembly (9). We also demonstrated that 6 human proteins, including homologs of our novel yeast targets, could be polyphosphorylated following the ectopic expression of *E. coli* PPK, the bacterial polyP polymerase, in HEK293T cells (9). Subsequent studies using protein microarrays identified an additional 8 human proteins as targets, in addition to 5 proteins that appear to bind polyP chains non-covalently (13). Altogether, work from our group and others suggests that lysine polyphosphorylation has the capacity to be a global protein modifier, akin to other lysine based modifications such as acetylation or ubiquitylation (8,14).

Our previous strategy to identify targets of lysine polyphosphorylation in yeast took advantage of a set of strains wherein each open reading frame is expressed as a fusion with the GFP epitope (15). Candidate fusions from this set were crossed into a wild-type or *vtc4*Δ mutant background prior to analysis by NuPAGE and immunoblotting with an antibody against GFP. In this assay, a *VTC4-*dependent electrophoretic shift signals a candidate polyphosphorylated protein. In our previous work (9), we prioritized 90 targets from 427 proteins containing putative PASK motifs, defined by a sliding window of 20 amino acids with 75 % D, E, S or K amino acids with at least 1 K. This set of 90 proteins was enriched for multiple or long PASK motifs containing many lysine residues. In an attempt to further define the landscape of polyphosphorylation in *S. cerevisiae*, we have now screened the remainder of the candidates present in this GFP collection (See Materials and Methods) using a similar strategy

## RESULTS & DISCUSSION

Using our previously defined screening approach, we have now uncovered an additional 8 candidate targets. As observed in our previous work, the shifts by polyP (i.e. in the presence of a functional *VTC4* gene) are dramatic and quantitative, with the whole population of target proteins being affected (**Figure 1**). We attribute this ‘all or nothing’ pattern to the high concentration of polyP in cells and in extracts following cell lysis, which could be quantitatively modifying proteins *in vitro. In vivo*, polyphosphorylation of any given target may be sub-stoichiometric. New polyphosphorylated targets uncovered in our screen include homologs of proteins implicated in human disease (**Figure 1, see below**). Despite the identification of proteins predicted to function in diverse processes, new targets display extensive physical and genetic interactions with previously identified polyphosphorylated proteins (**Figure 2**). Heh2 is an inner nuclear membrane protein involved in ensuring the quality control of nuclear pore complexes (16). Ygr237c and Kel3 are uncharacterized proteins with diverse physical and genetic interactions and putative cytoplasmic localization (9). Itc1 is a member of the Itc1-Isw2 chromatin remodeling complex involved in meiotic gene repression (17). Nop2 is an rRNA methyltransferase required for 27S rRNA processing and 60S ribosome biogenesis (18,19). While largely uncharacterized, Syh1 co-purifies with ribosomes (20), suggesting that it too functions in translation or ribosome assembly. *SYH1* also displays genetic interaction with *EBP2*, encoding a protein involved in 60S biogenesis. The identification of Nop2 and Syh1 are consistent with our previous observation that *vtc4*Δ cells show defects in polysome formation and the previous identification of negative genetic interactions between *vtc4*Δ mutants and deletions of genes impacting ribosome biogenesis (*RPP1A, MAK11*, and *RPC25*) and translation (*ELP3, ELP6*, and *MNR1*) (21,22). Our expanded target list also included a number of proteins associated with vacuolar biology. Prb1 encodes the yeast vacuolar proteinase B (23). Apl5 is involved in transporting protein cargo from the ER-golgi system to the vacuole (24). Since the vast majority of polyP localizes to the vacuole in *S. cerevisiae*, these proteins are ideal candidate effectors of polyP-mediated signaling. Therefore, we carried out further experiments to validate the polyphosphorylation of these new targets and to determine amino acid sequences required for the modification.

**Figure 1.**
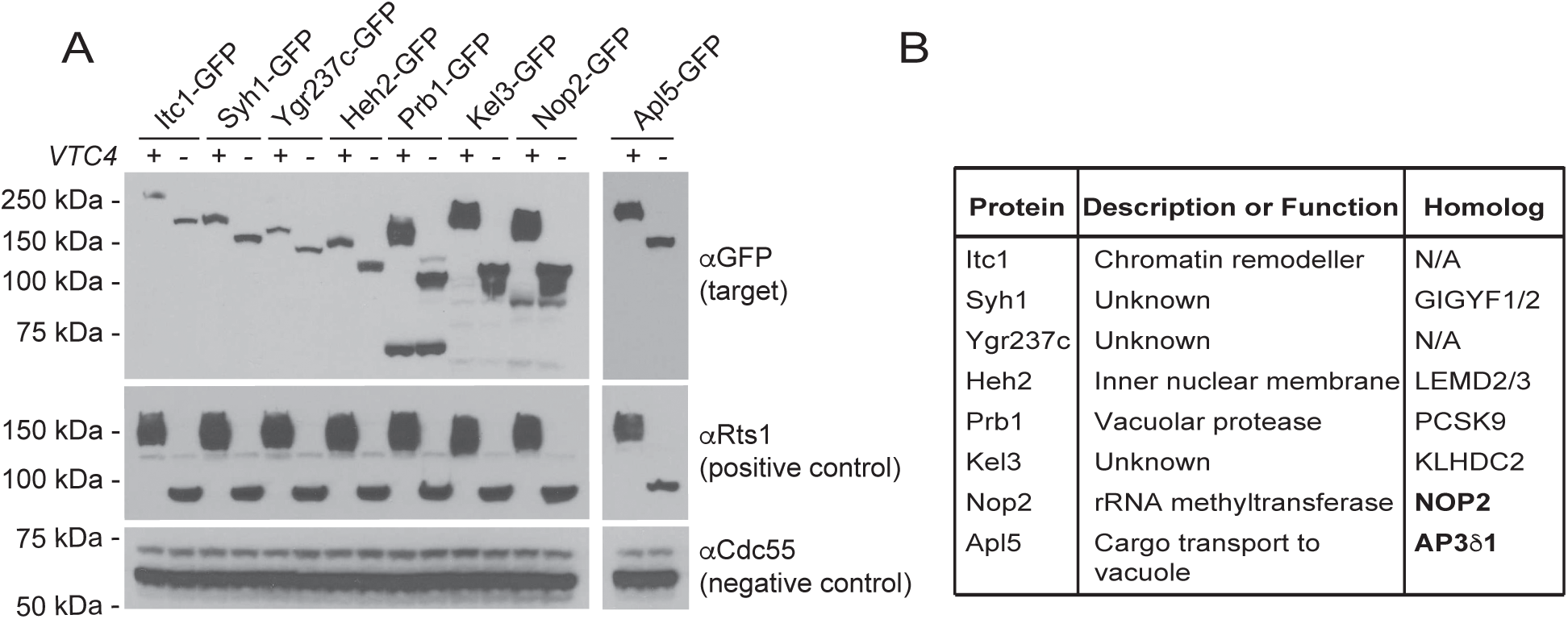
Identification of new polyphosphorylated targets in yeast. **A)** NuPAGE analysis of polyphosphorylated proteins. Paired GFP-tagged proteins were extracted from wild-type or *vtc4*Δ mutant cells using TCA lysis prior to NuPAGE analysis, western blotting to PVDF membrane and detection using the indicated antibodies. Rts1 serves as a positive control. Cdc55 serves as a negative control. **B)** Table of new polyphosphorylated targets uncovered in this study. Human homologs were determined using the Yeastmine tool. Bold indicates presence of a PASK sequence in human homolog (as defined by Bentley-DeSousa *et al.* 2018).

**Figure 2.**
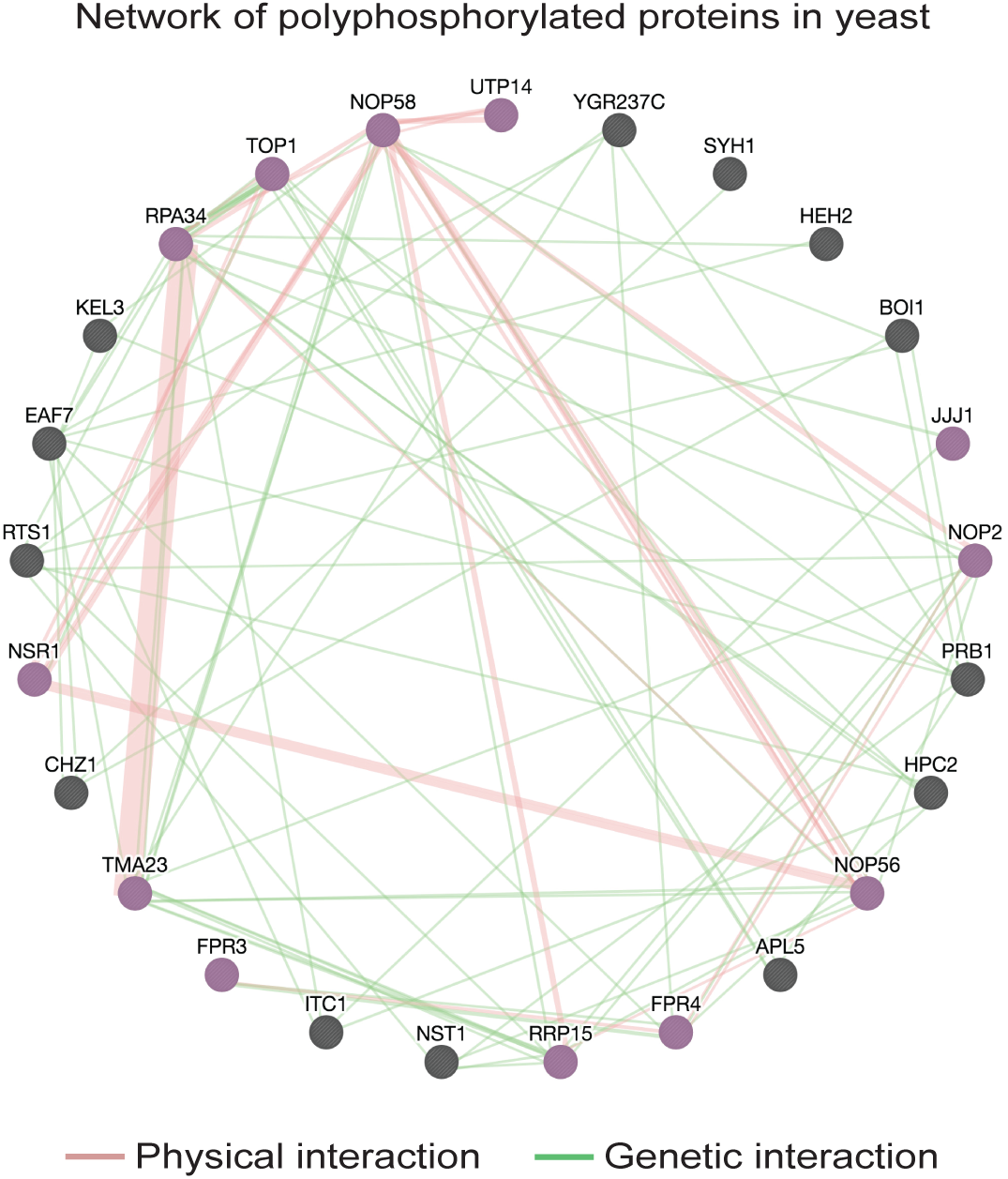
Polyphosphorylated targets are connected by genetic and physical interactions. Genetic and physical associations between new and previously identified targets were computed using the Genemania tool (see Materials and Methods). Each node (circle) represents an individual polyphosphorylated protein (n = 22). Physical and genetic interactions are indicated with green and pink lines respectively. Purple nodes indicate localization to the nucleolus (n= 12; FDR = 1.85 × 10^−7^).

We first sought to confirm that the electrophoretic shift detected for Prb1-GFP in the presence of *VTC4* was due to polyP. Since polyphosphorylation occurs non-enzymatically even in harsh conditions (7), we added synthetic polyP of 75 units in length to *vtc4*Δ whole-cell protein extracts generated via a trichloroacetic acid (TCA) lysis protocol. Addition of 5 mM of polyP conferred an electrophoretic shift to Prb1-GFP in *vtc4*Δ extracts that was equal to the shift observed in the wild-type control. In contrast, the same concentration of sodium phosphate had no impact on the mobility of Prb1-GFP, confirming that polyP is required for the modification (**Figure 3A**).

**Figure 3.**
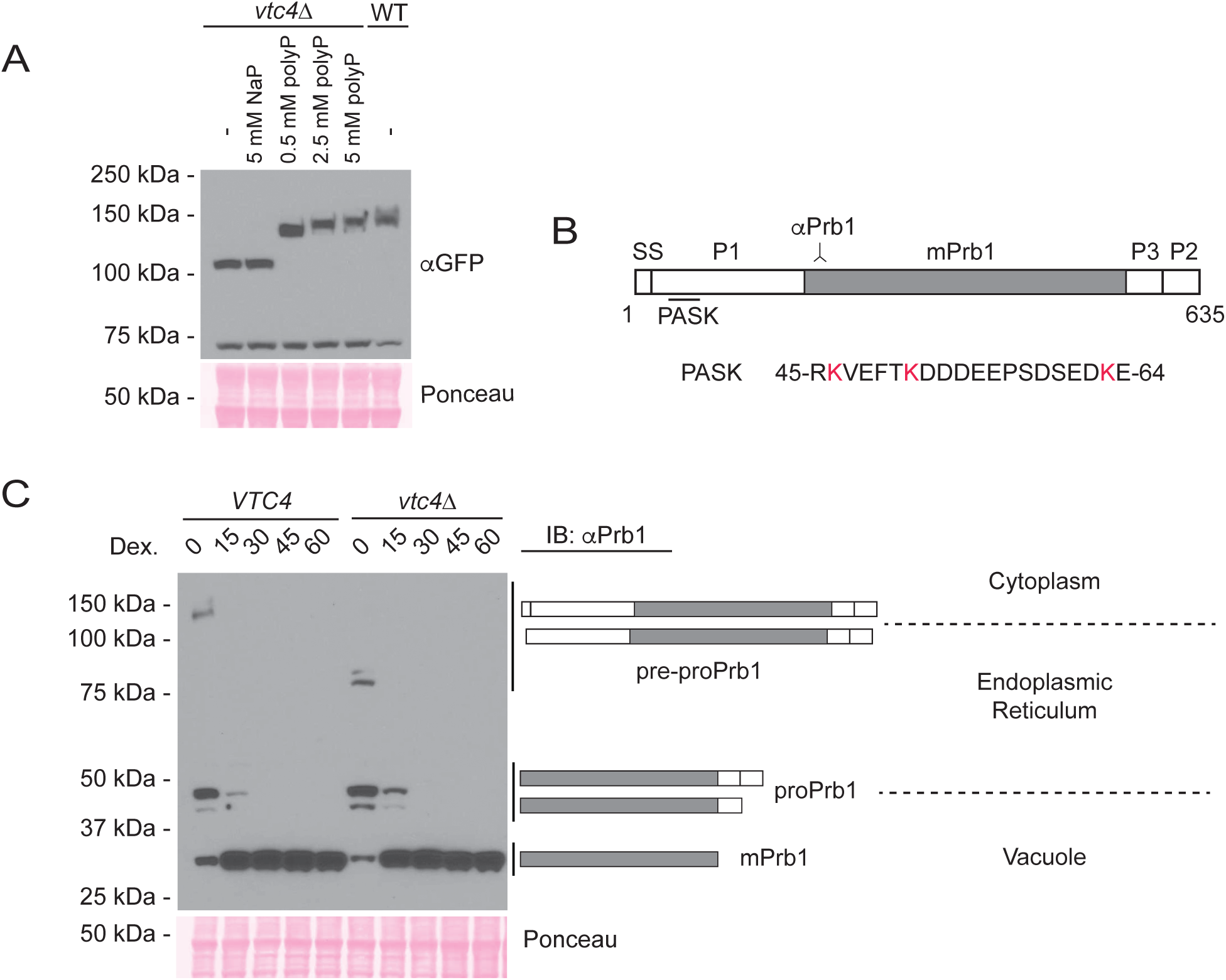
Analysis of Prb1 polyphosphorylation. **A)** Indicated concentrations of polyP or sodium phosphate control were added to *vtc4*Δ mutant cell extracts expressing GFP-tagged Prb1 and polyP-induced shifts were analyzed by western blotting with an antibody against GFP after NuPAGE analysis. **B)** Schematic of inactive pre-pro form of Prb1. The region recognized by the antibody is shown. Red lysines indicate candidate sites for polyP attachment. **C)** Prb1 was expressed under the GAL promoter (see Materials & Methods) for 2 hours prior to addition of glucose to stop transcription. Processing and polyphosphorylation of Prb1 at the indicated time points was monitored via NuPAGE analysis. The proposed schematic for Prb1 processing indicated on the right was adapted from that proposed in Mark *et al.* 2015 (27).

The Prb1 protease is a heavily processed protein that undergoes a series of glycosylations and programed cleavages to convert the zymogen precursor (pre-pro) form to the fully active enzyme (mPrb1) (25). Processing occurs in step with its transit through the ER-golgi system to the vacuole (**Figure 3B & C**). Since the C-terminal GFP tag of Prb1-GFP is predicted to be removed during its processing, **Figures 1A** and **3A** only confirmed polyphosphorylation of pre-proPrb1. In order to study polyphosphorylation of Prb1 in the context of its processing from zymogen to active enzyme, we used a previously described galactose overexpression system (26). Induction of *PRB1* transcription with the addition of galactose, followed by addition of glucose to turn off that transcription, allowed for the production of a population of native Prb1 whose processing could be monitored over time. Following analysis by NuPAGE and western blotting, we detected Prb1 with a previously published antibody that recognizes the first 14 amino acids of the fully processed and active (mPrb1) protein (**Figure 3B**) (25). We expect this antibody to detect all forms of Prb1. Only pre-proPrb1, but not any of the processed forms, exhibited an electrophoretic shift in *VTC4* strains relative to *vtc4*Δ mutants (**Figure 3C**). As a control, the previously identified target Rts1 (9) showed polyphosphorylation throughout the time course in *VTC4* cells (**Supporting Figure 1A**). These data suggest that the P1 segment of Prb1, which is removed by the Pep4 protease in the first cleavage event within the endoplasmic reticulum (25), is the modified species. Indeed, the predicted PASK motif is located in the P1 fragment, as indicated in **Figure 3B**. Importantly, analysis of these same samples via traditional SDS-PAGE, which does resolve polyP-induced shifts (9), showed identical migration of all species, consistent with the electrophoretic shift being due to polyphosphorylation (**Supporting Figure 1B**). In this analysis, we also observed slightly delayed processing of Prb1 in *vtc4*Δ mutants, although it is unclear if this is biologically meaningful (**Supporting Figure 1B**).

Polyphosphorylation of the P1 fragment may be required to ensure early processing steps occur in a manner consistent with the production of an active mPrb1 protein. In this capacity, polyP may function redundantly or in coordination with other PTMs, including glycosylations. Alternatively, polyP may function in a quality control capacity to regulate the function or degradation of Prb1 that is improperly processed by the ER-golgi system. A similar role has been suggested for the SCF^Saf1^ ubiquitin ligase, which ubiquitylates Prb1 but does not impact the bulk turnover of the protein population (26,27). The hypothesis that polyP modulates protein stability is consistent with its previously described role as a molecular chaperone in other biological systems (28).

Apl5, the second vacuole-related protein uncovered in our screen, functions as a member of the conserved AP3 complex. AP3 selectively transports protein cargo from the Golgi to lysosome-related organelles such as the yeast vacuole (29,30). As with Prb1-GFP, Apl5-GFP polyphosphorylation was rescued by the addition of polyP to *vtc4*Δ extracts in a concentration-dependent manner (**Figure 4A**). Other subunits of AP3 were not polyphosphorylated (**Figure 4B**), suggesting that Apl5 may be a focal point for AP3 regulation by polyP. As predicted, the PASK motif of Apl5 is required for the electrophoretic shift observed in *VTC4* strains (**Figure 4C**). This PASK motif is located in the ‘ear domain’ of the protein (**Figure 4D**), previously demonstrated to be required for interaction with AP3 regulators such as Vps41 (24,31). This interaction occurs when Vps41 is integrated into the HOPS complex, and is thought to be required for AP3 docking to the vacuole membrane, which allows proper delivery of vacuolar cargo (24). The other large subunit of the AP3 complex, Apl6, also has a PASK motif (9), although we did not detect reproducible Vtc4-dependent electrophoretic shifts of this protein on NuPAGE gels (**Figure 4B**). Notably, Apl6 only has 2 lysines in its PASK motif, compared to 12 for Apl5 (**Supporting Figure 2A**). Since we previously found that electrophoretic shifts due to polyphosphorylation scale in proportion to the number of lysines in the PASK motif (9), it is possible that Apl6 polyphosphorylation is not easily detected via this method.

**Figure 4.**
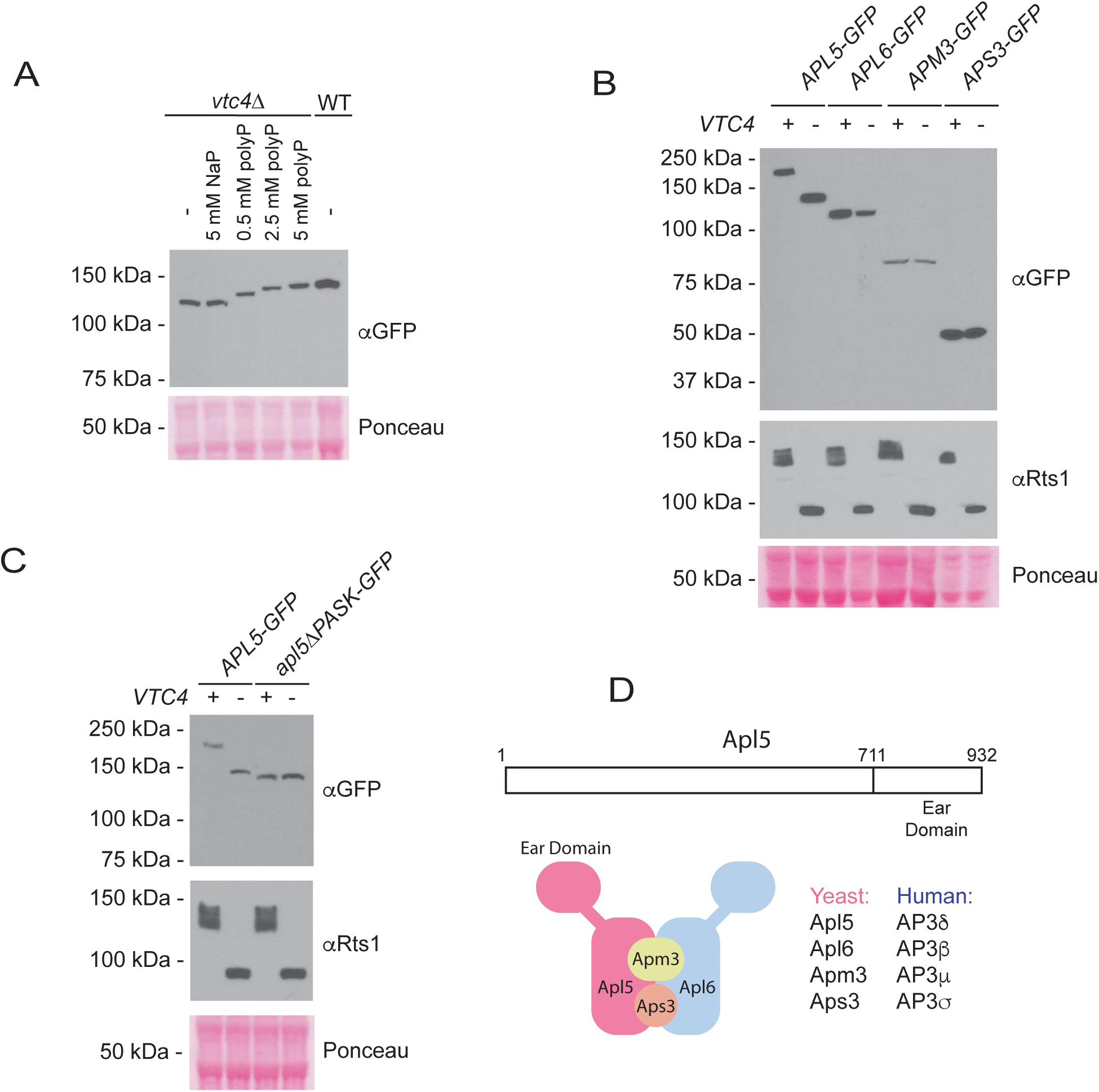
Analysis of Apl5 polyphosphorylation. **A)** Indicated concentrations of polyP or sodium phosphate control were added to *vtc4*Δ mutant cell extracts expressing GFP-tagged Apl5 and polyP-induced shifts were analyzed by western blotting with an antibody against GFP after NuPAGE analysis. **B)** The indicated components of the AP3 complex were expressed as GFP fusions in WT and *vtc4*Δ strains and analyzed by western blotting following NuPAGE analysis of protein extracts. **C)** The PASK motif of Apl5 is required for its polyphosphorylation. Protein extracts from the indicated strains were analyzed as described in B. **D)** Representation of the conserved Apl5 protein and the AP3 complex. The schematic of the AP3 complex is adapted from that proposed previously (29).

AP3 mutant strains were previously described as having low levels of polyP (32). AP3 could impact polyP synthesis by transporting proteins involved in polyP synthesis or storage to the vacuole (32–34). In the context of this model, polyphosphorylation of AP3 may serve as means of feedback to regulate AP3 function. Importantly, the impact of AP3 on polyP metabolism could be mediated by multiple cargos, as over 200 genes are known to impact polyP synthesis in *S. cerevisiae* (32). In humans, mutations in the AP3β and AP3δ subunits of the AP3 complex, homologs of Apl6 and Apl5 respectively (**Figure 4D**)(29), give rise to Hermansky-Pudlak Syndrome (HPS) subtypes 2 and 10 (35–37). HPS is a pleiotropic disease characterized by defects in the transport of proteins to lysosome related-organelles (38,39). Notably, a hallmark of HPS is bleeding diathesis and a failure to accumulate or retain polyP in the dense granules of platelets (38,39). PolyP has a well-characterized role in the blood coagulation cascade and works by stimulating activation of several clotting factors (14). AP3β1 AP3β2 and AP3δ also have PASK motifs as defined by our criteria (**Supporting Figure 2B**), and Azevedo et al. provided evidence that AP3β1 can be polyphosphorylated (13). Direct regulation by polyP may be a conserved feature of AP3 biology.

Altogether, our work expands the scope of polyphosphorylation in yeast and provides new avenues for exploring the functional consequences of this intriguing post-translational modification. Expanding the catalog of polyphosphorylated proteins in other eukaryotes will be a critical step in understanding the molecular and cellular functions of polyP in these systems.

## METHODS

### Yeast strains

All strains are in the S288C background and were generated using standard methods. Strains were verified using PCR to confirm the presence of the knockout cassette and the absence of the wild-type open reading frame. This analysis was also done for strains streaked or generated from strains taken from the GFP-tagged set. Strain confirmations were always done on colony-purified isolates, whose genotypes are described in **Supporting Table 1.** Details of individual strain constructions, including oligonucleotides used for tagging, deletion or confirmations, are available upon request. For the GAL shut-off experiment used to examine Prb1 processing, wild-type *GALpr-PRB1* and *GALpr-PRB1 vtc4*Δ mutant strains were grown in synthetic complete (SC) media with 2 % raffinose overnight. Cells were diluted to OD600 = 0.3 in SC media with 2 % raffinose and grown until OD600 = 0.6. Cultures were then supplemented with 2 % galactose for 30 minutes to induce Prb1 expression. Subsequently, glucose was added to a final concentration of 2 % and time points were taken every 15 minutes. For all other experiments, cells were grown in YPD media supplemented with 0.005 % adenine and tryptophan. All strains were grown at 30 °C.

### Screening protocol

Candidate target strains were recovered from the yeast GFP-tagged collection and the *vtc4*Δ and *ppn1*Δ mutations were crossed into the strains as previously described (9). Paired protein extracts were generated and analyzed using NuPAGE analysis following preparation of protein extracts via TCA lysis (see below). Rts1 was used as control for a polyphosphorylated protein for each sample analyzed. Candidates were confirmed in a secondary screen before streaking strains for single colonies and confirming the correct position of the GFP tag. As described throughout the results section, select hits were reconfirmed by re-tagging candidates in various genetic backgrounds and/or with the use of antibodies directed against native targets. A list of screened targets and their predicted PASK motifs is included in **Supporting Table 1.**

### Protein analysis

Protein extraction was done using a TCA lysis method that has been described previously (9). A similar protocol is reiterated here. Briefly, 3-6 OD600 equivalents of yeast cells were lysed in 300 μL of 20 % TCA with 100 μL of acid-washed glass beads. Supernatant was recovered and beads were washed with 300 μL of 5 % TCA and combined with the first supernatant. Samples were centrifuged at 16,000 × g for 4 minutes at 4 °C. Pellets were resuspended in 100 μL SDS-PAGE sample buffer (0.8 mL 3X SDS PAGE sample buffer, 100 μL 1.5 M Tris-HCl pH 8.8, 100 μL 1 M DTT). Samples were boiled for 5 minutes and centrifuged again at 16,000 × g for 4 minutes. Supernatant was removed for immediate analysis or stored at −80 °C until use. Typically, 10-20 μL of extract was loaded for each lane. We have observed that protein samples tend to lose their polyphosphorylation after multiple freeze thaw cycles. We recommend same-day analysis of extracted proteins.

### Western blotting

Buffers used for NuPAGE and SDS-PAGE analysis were described previously (9). SDS-PAGE gels used for this work were 10 % final concentration made with 30 % acrylamide/bis solution at a ratio of 37.5:1 (BioRad 1610158). Please note that SDS-PAGE analysis does not resolve polyP-induced shifts. PVDF membrane (Biorad 162-0177) was used for Western blotting of separated protein samples. Primary and secondary antibodies used for immunoblotting, including dilution information for each antibody, is described in **Supporting Table 2**. Washed blots were exposed to ECL from Millipore (Sigma WBKLS0500 or WBLUF0500). Ponceau S (Sigma P-3504) was used to verify equal loading across the blots and to monitor the quality of transfer for each blot.

### Homologs and Network analysis

Homologs were determined via the Yeastmine tool (40). Genetic networks were generated using Genemania (genemania.org; June 12, 2019) (41). Gene attributes were set to zero, only genetic and physical interactions were computed and shown, all other settings were left as default. FDR displayed was as calculated for nucleolar localization by the Genemania tool.

### In vitro polyphosphorylation assays

*In vitro* polyphosphorylation assays were carried out by mixing the indicated concentrations of polyP with TCA-style protein preps, incubation at room temperature for 30 minutes before reboiling of reactions and NuPAGE analysis, as described above. PolyP chains used for this analysis are an average of 75 residues in length and were purchased from Kerafast (Medium chain, EUI005).

## ACKNOWLEDGEMENTS

We thank the members of the Downey lab for critical reading of the manuscript and the Dr. Elizabeth Jones’ lab for the anti-Prb1 antibody. ABD was supported by an Ontario Graduate Scholarship. This work was funded by a Canadian Institutes of Health Research Project Grant to MD (PJT-148722). We also acknowledge funding from a Rare Disease Foundation Microgrant (grant number 1937).

## CONFLICTS OF INTEREST

The authors declare no conflicts of interest.

---

**Supplementary Figure 1.**
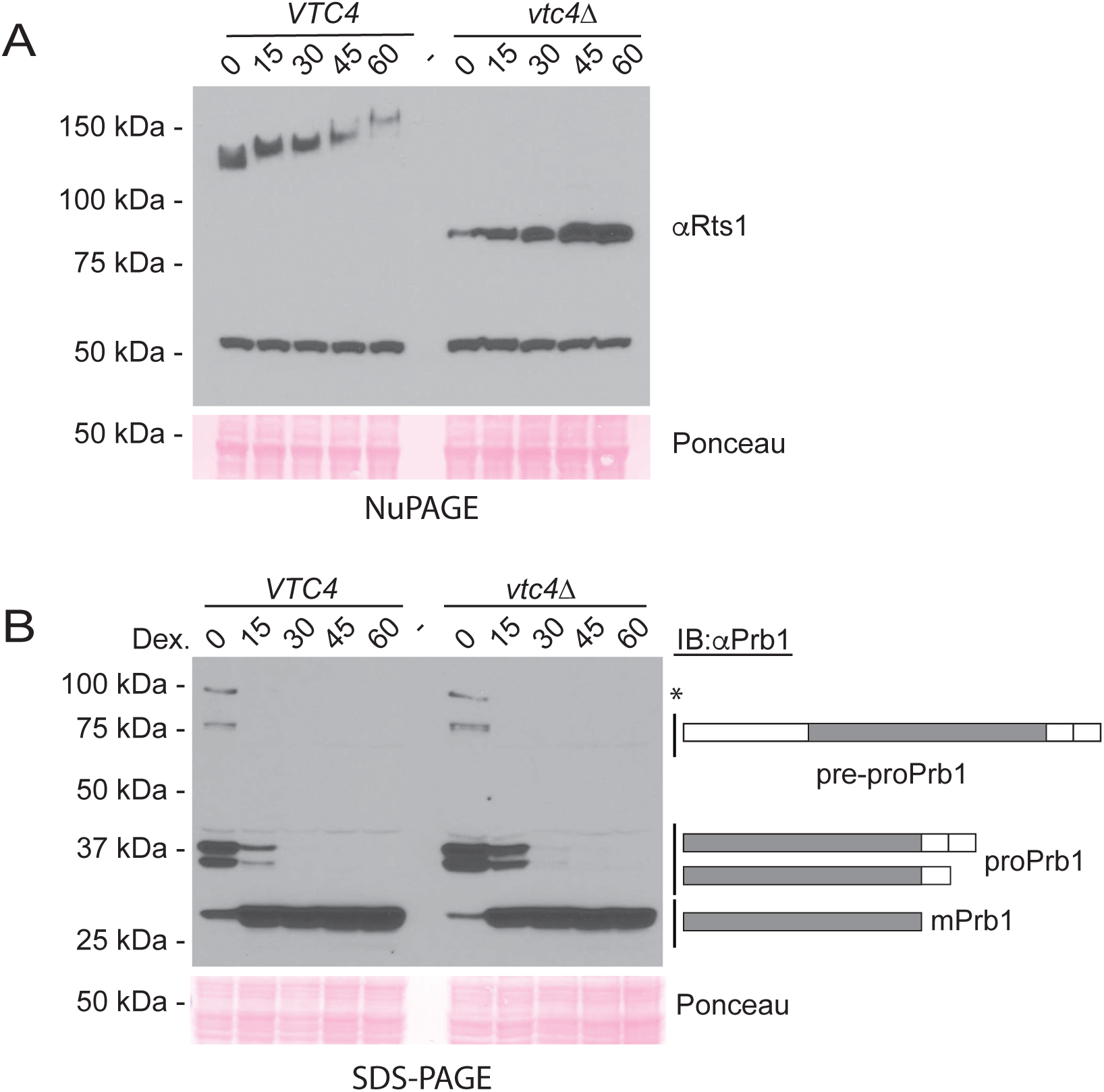
Analysis of Prb1 processing and polyphosphorylation. **A)** Polyphosphorylation of Rts1 was monitored during the experiment presented in Figure 4C using an antibody against Rts1. **B)** Processing of pre-pro Prb1 was monitored for the experiment presented in Figure 4C using SDS-PAGE and immunoblotting with an antibody against Prb1. Note: SDS-PAGE does not resolve shifts due to lysine polyphosphorylation. * may represent a previously uncharacterized form of Prb1. **Relates to Figure 3.**

**Supplementary Figure 2.**
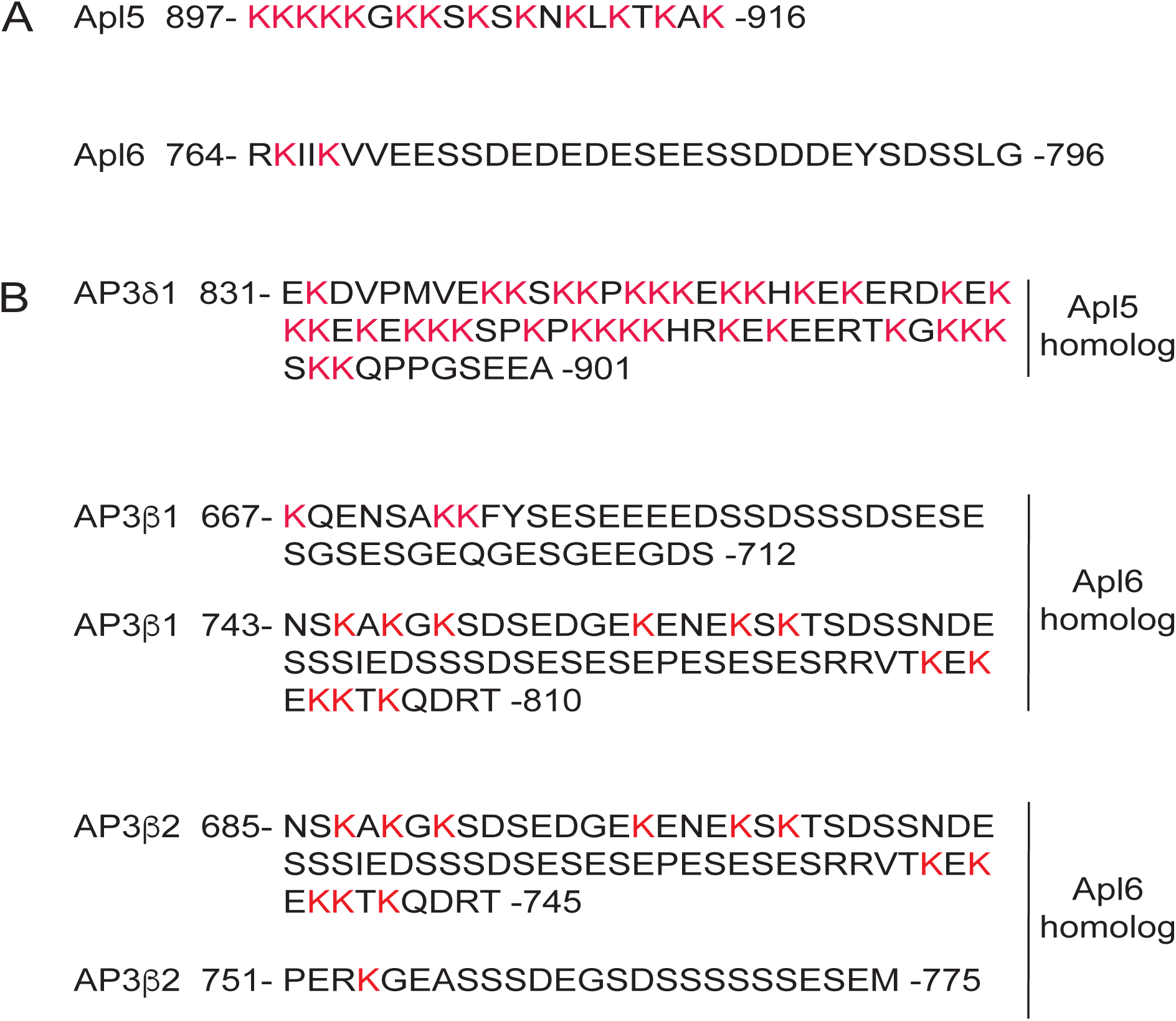
PASK sequences within the AP3 complex. **A)** PASK sequences of Apl5 and Apl6. Numbers indicate amino acid positions. **B)** PASK sequences of AP3β and AP3δ. Numbers indicate amino acid positions. **Relates to Figure 4.**

## SUPPORTING TABLES: (Provided in a single excel sheet)

**Supporting Table 1** - PASK containing proteins screened for polyphosphorylation in this study.

**Supporting Table 2** - Yeast strains used in this study.

**Supporting Table 3** - Antibodies used in this study.

## Notes

#### Summary of Updates

I have updated the supplemental table 1 to provide an indicate of some strains that were not ultimately assayed for polyphosphorylated because they were not recovered from the SGA-style crosses.

